# Tartrolon sensing and detoxification by the *Listeria monocytogenes timABR* resistance operon

**DOI:** 10.1101/2023.07.27.550809

**Authors:** Tim Engelgeh, Jennifer Herrmann, Rolf Jansen, Rolf Müller, Sven Halbedel

## Abstract

*Listeria monocytogenes* is a foodborne bacterium that naturally occurs in the soil. Originating from there, it contaminates crops and infects farm animals and their consumption by humans may lead to listeriosis, a systemic life-threatening infectious disease. The adaptation of *L. monocytogenes* to such contrastive habitats is reflected by the presence of virulence genes for host infection and other genes for survival under environmental conditions. Among the latter are ABC transporters for excretion of antibiotics produced by environmental competitors, however, most of these transporters have not been characterized. Here, we generated a collection of promoter-*lacZ* fusions for genes encoding ABC-type drug transporters of *L. monocytogenes* and screened this reporter strain collection for induction using a library of natural compounds produced by various environmental microorganisms. We found that the *timABR* locus (*lmo1964-lmo1962*) was induced by the macrodiolide antibiotic tartrolon B, which is synthesized by the soil myxobacterium *Sorangium cellulosum*. Tartrolon B resistance of *L. monocytogenes* was dependent on *timAB*, encoding the ATPase and the permease component of a novel ABC transporter. Moreover, transplantation of *timAB* was sufficient to confer tartrolon B resistance to *Bacillus subtilis*. Expression of the *timABR* locus was found to be auto-repressed by the TimR repressor, whose repressing activity was lost in the presence of tartrolon B. We also demonstrate that tartrolon sensitivity was suppressed by high external potassium concentrations, suggesting that tartrolon acts as potassium ionophore. Our results help to map the ecological interactions of an important human pathogen with its co-residing species within their joint natural reservoir.

**Importance:** *Listeria monocytogenes* is an environmental bacterium that can cause listeriosis in humans when ingested. The genome of this bacterium contains many genes encoding putative ATP binding cassette transporters with unknown function. Presumably, these transporters serve the excretion of antimicrobial compounds produced by co-residing species competing with *L. monocytogenes* for nutrients and habitats. We here fused the *lacZ* reporter gene to the promoters of these transporter genes and screened a natural compound library for activating substances. We discovered that tartrolon B activates expression of the *L. monocytogenes timABR* operon, encoding the TimAB exporter and the TimR repressor. Tartrolon B is an antibiotic produced by the soil-dwelling myxobacterium *Sorangium cellulosum*. We elucidate how the *timABR* genes mediate sensing and detoxification of this antibiotic. This represents the first known mechanism of tartrolon resistance and may help to better define the natural reservoir of *L. monocytogenes*.

## Introduction

*Listeria monocytogenes* is the causative agent of listeriosis, a dreaded foodborne infection in humans and farm animals. The bacterium belongs to the *Bacillota* phylum (formerly known as firmicutes) and can be found in different environmental habitats including the soil, surface waters, plant surfaces and the gut of various animals (1-4). Contamination of crops and colonization of farm animals is the gateway for *L. monocytogenes* to be carried over into food production facilities, where the bacterium may colonize equipment, forms biofilms and finally contaminates food items (5). After ingestion, *L. monocytogenes* is capable to breach the gut epithelium and to enter the blood stream, but this early stage of the infection is normally brought under control by the immune system (6). However, the infection may spread to secondary organs in vulnerable risk groups, which then results in invasive listeriosis associated with high case fatality rates (7, 8). A combination of virulence factors and their regulatory elements orchestrates survival of *L. monocytogenes* in the infected host, while general and specific stress response genes are induced to cope with harmful conditions that prevail in environmental habitats (5, 9). Among the latter are multidrug resistance (MDR) transporters excreting toxic compounds such as bile or ethidium bromide as well as antibiotics (10).

*L. monocytogenes* MDR transporters with known substrate specificities belong to at least three different classes: The major facilitator superfamily (MFS) transporters such as MdrL (for export of macrolides and ethidium bromide) and Lde (fluoroquinolones, ethidium bromide) use the proton motive force (PMF) as energy source to drive compound extrusion (10-12). Likewise, QacH, a representative of the small MDR (SMR) transporters, is PMF-dependent; it transports quaternary ammonium compounds out of the cell (13). The ATP binding cassette (ABC) type MDR transporters constitute a third but different class as they directly use ATP hydrolysis for compound extrusion. Transporters of this class consist of an ATPase coupled to a permease unit either separated on two polypeptides or fused in a single protein (10). Even though systems of this type are frequently regarded as MDR transporters, it often remains questionable whether they really aid in the transport of different compound classes or whether they are rather SDR (single drug resistance) transporters (14, 15). Several *L. monocytogenes* ABC transporters are known to contribute to toxic compound excretion such as BilEAB (bile), AnrAB (bacitracin, nisin, cephalosporins) or LieAB (aurantimycin) (16-20), and many more ABC-type transporters possibly related to compound export are encoded in the genome, however, most of them have not been characterized. We originally identified the *lieAB* genes while searching for target genes of the LftR transcriptional repressor (18, 21). LftR is a PadR-type repressor that silences expression of its target genes (including *lieAB*) by binding as a dimer to the partially palindromic GTAWTAC-N_3_-ATAC operator sequence found in LftR-regulated promoters (19, 22). Repression of *lieAB* transcription is relieved upon aurantimycin exposure in a process that requires LftS, a co-regulator of LftR; and induction of LieAB production then confers aurantimycin resistance (18, 19). Whether aurantimycin is sensed by LftR directly or in complex with LftS is currently not clear (19). Aurantimycin is a cyclic hexadepsipeptide, produced by soil-dwelling *Streptomyces aurantiacus* which forms pores in lipid membranes (23, 24), whereas bacitracin (the substrate of AnrAB) is a non-ribosomally synthesized macrocyclic peptide produced by several *Bacillus licheniformis* strains (25, 26) and nisin (another AnrAB substrate) is a gene-encoded and lanthionine ring-containing antimicrobial peptide produced by *Lactococcus lactis* (27). Like aurantimycin-producing *S. aurantiacus* that can be isolated from soil (28, 29), *B. licheniformis* (bacitracin producer) and *L. lactis* (nisin producer) are found in similar environmental niches (soil, plant surfaces) as *L. monocytogenes* (30, 31). Moreover, expression of the *anrAB* ABC transporter genes is also induced by the same substances that are exported by their gene products (20), as reported for the *lieAB* genes (18). However, control of *anrAB* expression is mediated by the VirRS two component system in conjunction with VirAB, a compound sensing ABC transporter (20) rather than by a transcriptional repressor.

We here describe the discovery of a novel *L. monocytogenes* ABC transporter, named TimAB. We show that TimAB contributes to the resistance against tartrolon antibiotics, which represent another group of antimicrobial compounds produced by soil-dwelling microorganisms. Furthermore, we show how TimAB production is controlled by the tartrolon-responsive transcriptional regulator TimR. Our results illustrate the tight exposure of an important human pathogen to a diverse spectrum of microbial competitors in its natural reservoir.

## Results

### Weakly transcribed MDR transporter genes in *L. monocytogenes* EGD-e

174 genes belonging to 104 putative or known ABC transporters are currently listed for *L. monocytogenes* EGD-e in the TransportDB database (32, 33). Among them, there are 23 uncharacterized ABC transporters either directly classified as MDR transporters or as being potentially involved in excretion of antibiotics (Tab. 1). Previously published RNA-Seq data from our lab (18) indicated that most of these transporter genes are not expressed under standard laboratory conditions (Tab. 1). Interestingly, some of the transporter genes are arranged in operons together with genes that are annotated as *gntR-, marR-, mngR-, lytR*- or *tetR-*like transcriptional regulators (Tab. 1), suggesting transcriptional induction during specific conditions. Most likely, these transcriptional regulators respond to low molecular weight ligands such as metabolic intermediates or toxic compounds including antibiotics (34-36). None of these 23 ABC transporters is associated with an extracellular substrate binding protein, further suggesting that they all may act as compound exporters. In order to identify conditions that lead to transcriptional induction of these uncharacterized MDR transporter genes or operons, we fused their 18 promoters to *lacZ* and inserted these promoter-*lacZ* fusions into the chromosome of strain EGD-e. These strains were then grown in BHI broth at 37°C to mid-logarithmic growth phase and β-galactosidase activity was determined. Promoter activity ranged between 14±3 miller units (MU) for P*_lmo0741_* and 356±64 MU for P*_lmo2215_* (Fig. 1). Median activity of the 18 tested promoters was 67 MU, which compares to the low activity of the P*_lieAB_* promoter when repressed by LftR (45±8 MU) and which is only somewhat higher than seen in a strain carrying the promoterless *lacZ* gene (13±3 MU). In contrast, relief of LftR repression of the P*_lieAB_* promoter in a Δ*lftR* background leads to a β-galactosidase activity as high as 8513±1508 MU (Fig. 1), which is in good agreement with previous results (18, 19). This shows that most of the MDR transporter genes are poorly expressed under standard growth conditions.

**Table 1:** ABC transporter genes in *L. monocyctogenes* EGD-e with possible functions in compound excretion (according to http://www.membranetransport.org).

| ATPase | permease | operon<sup>1</sup> | TPM<sup>2</sup> |
| --- | --- | --- | --- |
|  | *lmo0107\** | *lmo0108 lmo0107* | 2 |
|  | *lmo0108\** |  | 2 |
| *lmo0194* | *lmo0195* | *lmo0193 lmo0194 lmo0195* | 50 |
|  | *lmo0607\** | <u>*lmo0606*</u> *lmo0607 lmo0608* | 51 |
|  | *lmo0608\** |  | 66 |
| *lmo0667* | *lmo0668* | *lmo0667 lmo0668<sup>#</sup>* | 60 |
| *lmo0742* | *lmo0743* | <u>*lmo0741*</u> *lmo0742 lmo0743 lmo0744 lmo0745* | 5 |
|  | *lmo0744\** |  | 2 |
|  | *lmo0925\** | *lmo0923 lmo0924 lmo0925 <u>lmo0926</u>* | 30 |
| *lmo0986* | *lmo0987* | <u>*lmo0984*</u> *lmo0985 lmo0986 lmo0987<sup>#</sup>* | 4 |
| *lmo1636* | *lmo1637* | *lmo1635 lmo1636 lmo1637* | 178 |
|  | *lmo1651\** | *lmo1652 lmo1651 lmo1650 lmo1649 lmo1648* | 28 |
|  | *lmo1652\** |  | 23 |
| *lmo1724* | *lmo1723* | <u>*lmo1725*</u> *lmo1724 lmo1723* | 22 |
| *timA* | *timB* | *lmo1964 (timA) lmo1963 (timB) <u>lmo1962 (timR)</u>* | 15 |
| *lmo2215* | *lmo2214* | *lmo2215 lmo2214* | 97 |
| *lmo2227* | *lmo2226* | *lmo2228 lmo2227 lmo2226* | 14 |
| *lmo2240* | *lmo2239* | <u>*lmo2241*</u> *lmo2240 lmo2239* | 70 |
| *lmo2372* | *lmo2371* | *lmo2371 lmo2372* |  |
|  | *lmo2745\** | monocistronic | 50 |
|  | *lmo2751\** | *lmo2751 lmo2752* | 38 |
|  | *lmo2752\** |  | 46 |
| *eslA* | *eslB* | *lmo2769 (eslA) lmo2768 (eslB) lmo2767 (eslC) <u>lmo2766 (eslR)</u>* | 30 |

<sup>1</sup> operons according to Toledo-Arana *et al.* (76), possible transcriptional regulators are underlined.
<sup>2</sup> TPM - transcripts per million as an estimate for gene expression according to previously published RNA-Seq data of *L. monocytogenes* wild type strain EGD-e during exponential growth, TPM values for the strictly repressed *lieAB* genes were ~10 (18).
\* contains both an ATPase and a permease domain in one polypeptide.
<sup>#</sup> not detected by Toledo-Arana *et al.* (76), but genetic arrangement suggests organization as an operon.

**Table 2:** Resistance of selected *Listeria* species against tartrolon B.

| species | group | timABR | MIC tartrolon B (µg/ml)<sup>1</sup> |
| --- | --- | --- | --- |
| *L. monocytogenes* | *Listeria sensu stricto* | + | 1.3±0.6 |
| *L. innocua* | *Listeria sensu stricto* | + | 6.6±2.3 |
| *L. ivanovii* | *Listeria sensu stricto* | + | 0.7±0.3 |
| *L. seeligeri* | *Listeria sensu stricto* | + | 1.3±0.6 |
| *L. aquatica* | *Mesolisteria* | + | >16 |
| *L. fleischmannii* | *Mesolisteria* | + | >16 |
| *L. floridensis* | *Mesolisteria* | + | 2.3±1.5 |
| *L. grayi* | *Murraya* | - | 0.004±0.003 |

**Figure 1:**
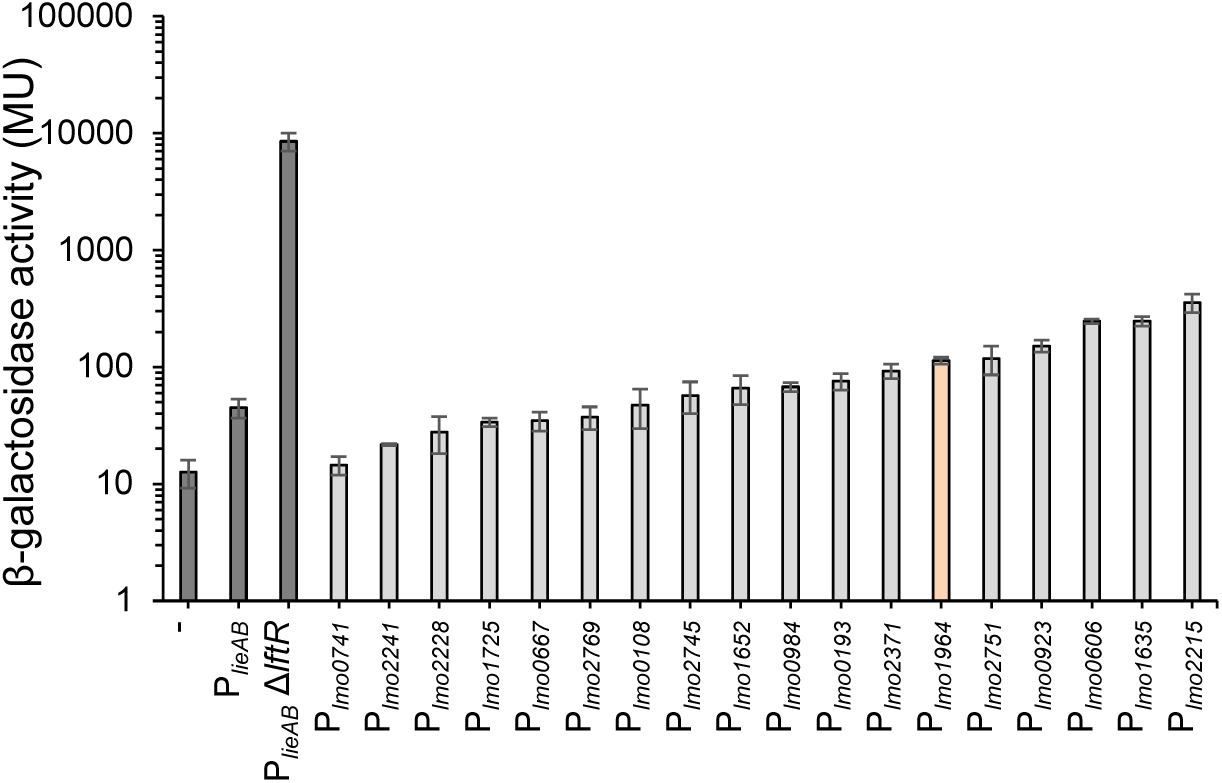
Promoters driving expression of potential MDR ABC transporter genes in *L. monocyctogenes*. Background activity of MDR ABC transporter gene promoters. Measurements of β-galactosidase activity in *L. monocytogenes* strains expressing promoter-*lacZ* fusions. Measurements were performed on strains grown in BHI broth at 37°C until midlogarithmic growth phase. Average values and standard deviations were calculated from three independent repetitions. Strains carrying a P*_lieAB_* promoter *lacZ* fusion either in wild type (LMSH5) or the Δ*lftR* background (LMSH98) or a promoter-less *lacZ* gene (LMSH16) were used as controls.

### Tartrolons induce expression of the *tim* operon

A natural compound collection was used to identify substances that induce the expression of the 18 ABC transporter gene promoters in a drop diffusion assay. This set as part of the DZIF (German Centre for Infection Research) natural product collection (37) contained 681 secondary metabolites purified from myxobacteria (253), streptomycetes (340) and fungi (88). The activity of the promoter of the *lmo1964-lmo1963-lmo1962* operon (Fig. 2) was found to be induced by tartrolons (Fig. 3A). This operon encodes the ATPase (*lmo1964*) and permease (*lmo1963*) subunits of an uncharacterized ABC transporter and a gene encoding a TetR-type transcriptional repressor (*lmo1962*, Fig. 2). For confirmation, promoter induction was tested with tartrolon A and B preparations, whose purity was separately validated using liquid chromatography - high resolution mass spectrometry (LC-hrMS, Fig. S1), and induction was observed with tartrolon B and – to a lesser extent – with tartrolon A as concluded from the appearance of a blue ring that surrounds the compound application site in the agar based diffusion assay with strain LMTE19 (*attB::*P*_lmo1964_- lacZ*) as the reporter strain. Both compounds generated zones of growth inhibition indicating toxicity (Fig. 3A). Tartrolons A and B are macrodiolide antibiotics isolated from *Sorangium cellulosum,* a soil-dwelling myxobacterium (38, 39). They consist of a 42-membered macrocyclic ring with four free hydroxyl groups in tartrolon A. In tartrolon B, however, these hydroxyl group oxygens are covalently bound to a single boron atom (38) (Fig. 3B). Tartrolons inhibit growth of different Gram-positive bacteria and block DNA, RNA and protein biosynthesis of *Staphylococcus aureus* (39). Tartrolons are structurally similar to boromycin (Fig. 3B), another boron containing macrocyclic antibiotic (40), however, boromycin did not induce the P*_lmo1964_* promoter (Fig. 3A).

**Figure 2:**
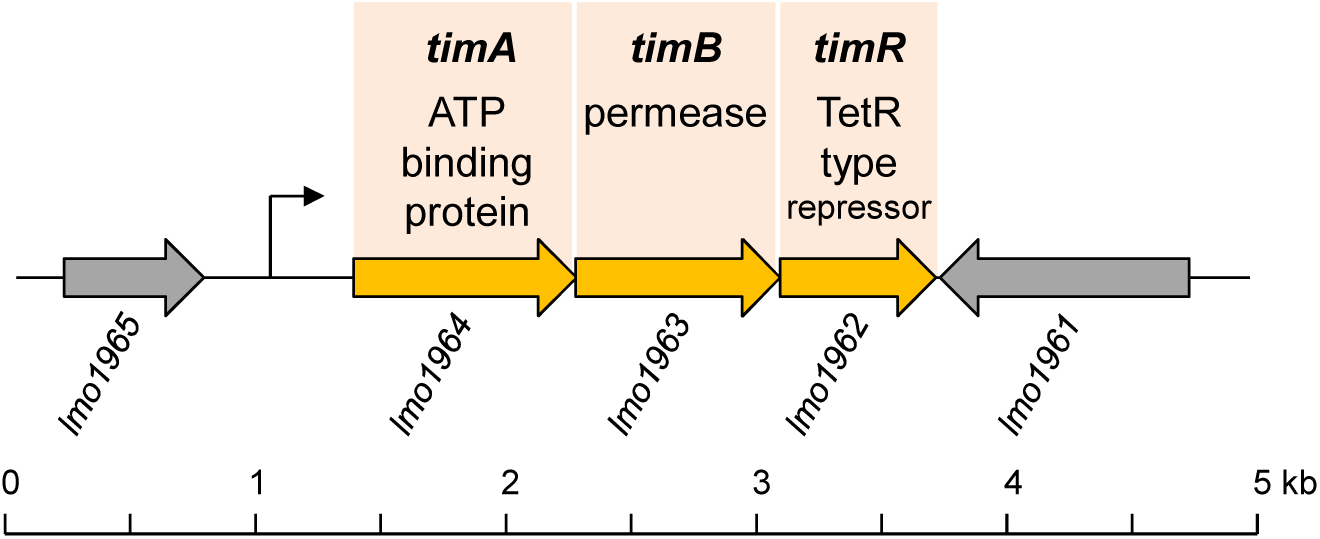
**Schematic illustration of the *timABR* ABC transporter locus of *L. monocytogenes* EGD-e.**

**Figure 3:**
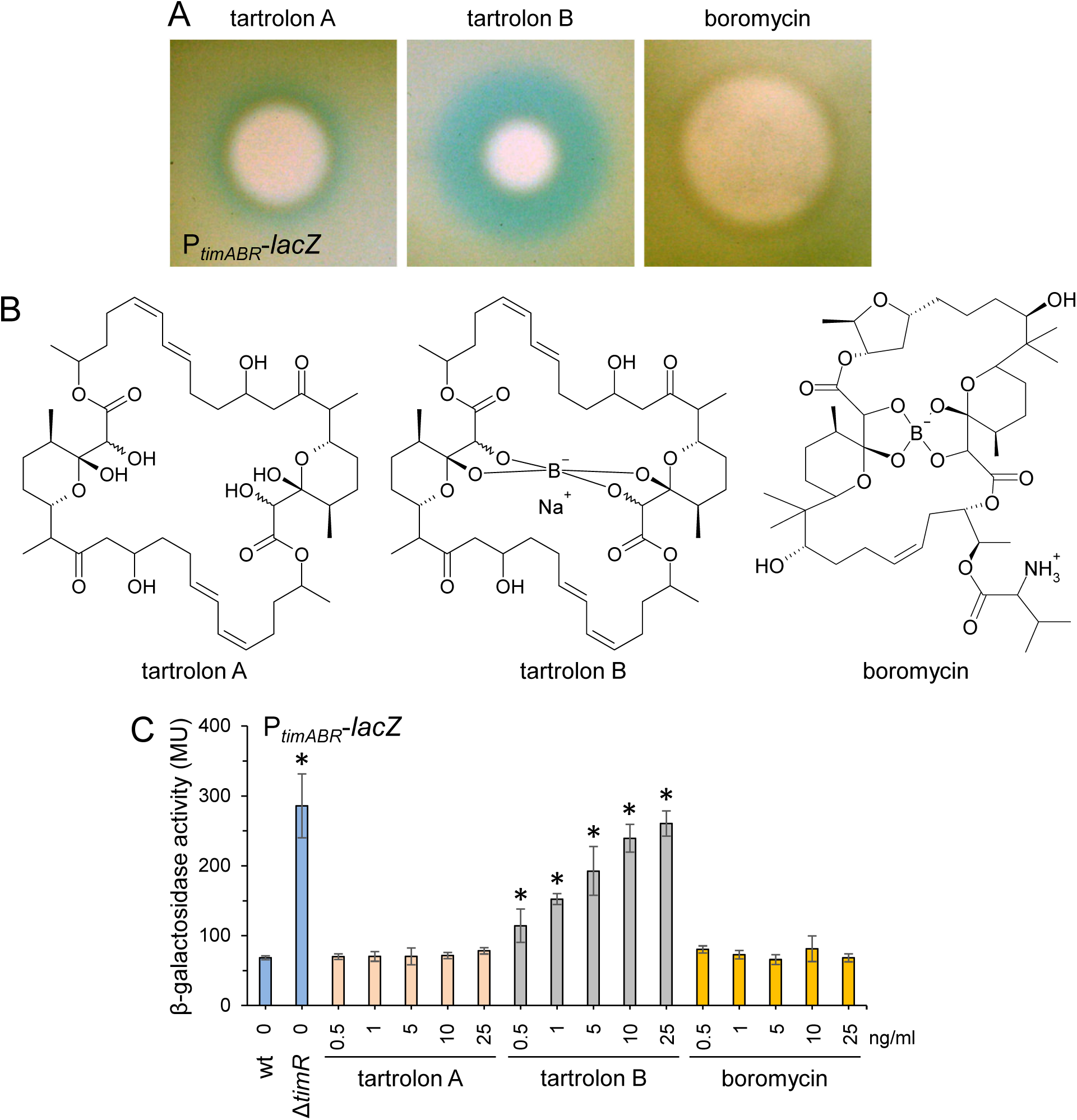
**Induction of the P*_timABR_* promoter by tartrolons A and B and boromycin.** (A) Induction of the P*_timABR_* promoter in an agar-based assay. Strain LMTE19 (P*_timABR_-lacZ*) was included in X-Gal containing BHI agar and 2 µl of compound solutions (2.5 µg/µl each) was spotted on top. The images were taken after overnight incubation at 37°C (B) Chemical structures of tartrolon A, tartrolon B and boromycin. (C) Concentration-dependent induction of the P*_timABR_* promoter by tartrolons and boromycin. LMTE19 (P*_timABR_-lacZ*) was grown in BHI broth containing increasing compound concentrations to mid-logarithmic growth phase at 25°C and β-galactosidase activity was determined. Strain LMTE50 (Δ*timR* P*_timABR_-lacZ*) was included for comparison. Average values and standard deviations were calculated from three independent repetitions. Asterisks mark statistically significant differences compared to LMTE19 without tartrolon (labelled “wt”, *P*<0.05, *t*-test with Bonferroni-Holm correction).

To further support the finding of tartrolon-specific induction of the P*_lmo1964_* promoter, β- galactosidase activity was determined in strain LMTE19 in the presence of increasing compound concentrations. While no induction was observed with tartrolon A and boromycin, tartrolon B induced this promoter in a concentration-dependent manner almost to the same degree as observed when the *lmo1962* repressor genes was deleted (Fig. 3C). Thus, tartrolon B is a specific inducer of the *L. monocytogenes* P*_lmo1964_* promoter.

### The *timABR* genes confer tartrolon resistance

Induction of P*_lmo1964_* by growth-inhibiting tartrolons raised the possibility that the potential MDR transporter encoded by the *lmo1964-lmo1963* genes could mediate resistance of *L. monocytogenes* against tartrolons. To test this hypothesis, a deletion mutant lacking the *lmo1964-lmo1963* genes was generated. Resistance of this mutant against tartrolon B was almost 200-fold reduced (minimal inhibitory concentration, MIC: 0.009±0.005 µg/ml) compared to wild type (MIC: 1.7±0.6 µg/ml), and a similar, but less pronounced effect was observed for tartrolon A (Fig. 4A). Reintroduction of an IPTG-inducible copy of the *lmo1964-lmo1963* genes into the Δ*lmo1964-lmo1963* mutant generated a strain that showed tartrolon A and B resistance similar to the Δ*lmo1964-lmo1963* mutant in the absence of IPTG and wildtype-like tartrolon resistance when IPTG was present (Fig. 4A). In agreement with its anticipated function as a repressor, deletion of *lmo1962* further increased tartrolon B resistance (MIC: 4±0 µg/ml) and reintroduction of the gene complemented this effect. In contrast, resistance against tartrolon A was not affected by the *lmo1962* deletion (Fig. 4A). Since the *lmo1964-lmo1962* operon can be induced by tartrolons and confers tartrolon resistance, we propose to rename these three genes as *timA* (*lmo1964*), *timB* (*lmo1963*) and *timR* (*lmo1962*, <u>t</u>artrolon <u>i</u>nducible <u>M</u>DR transporter).

**Figure 4:**
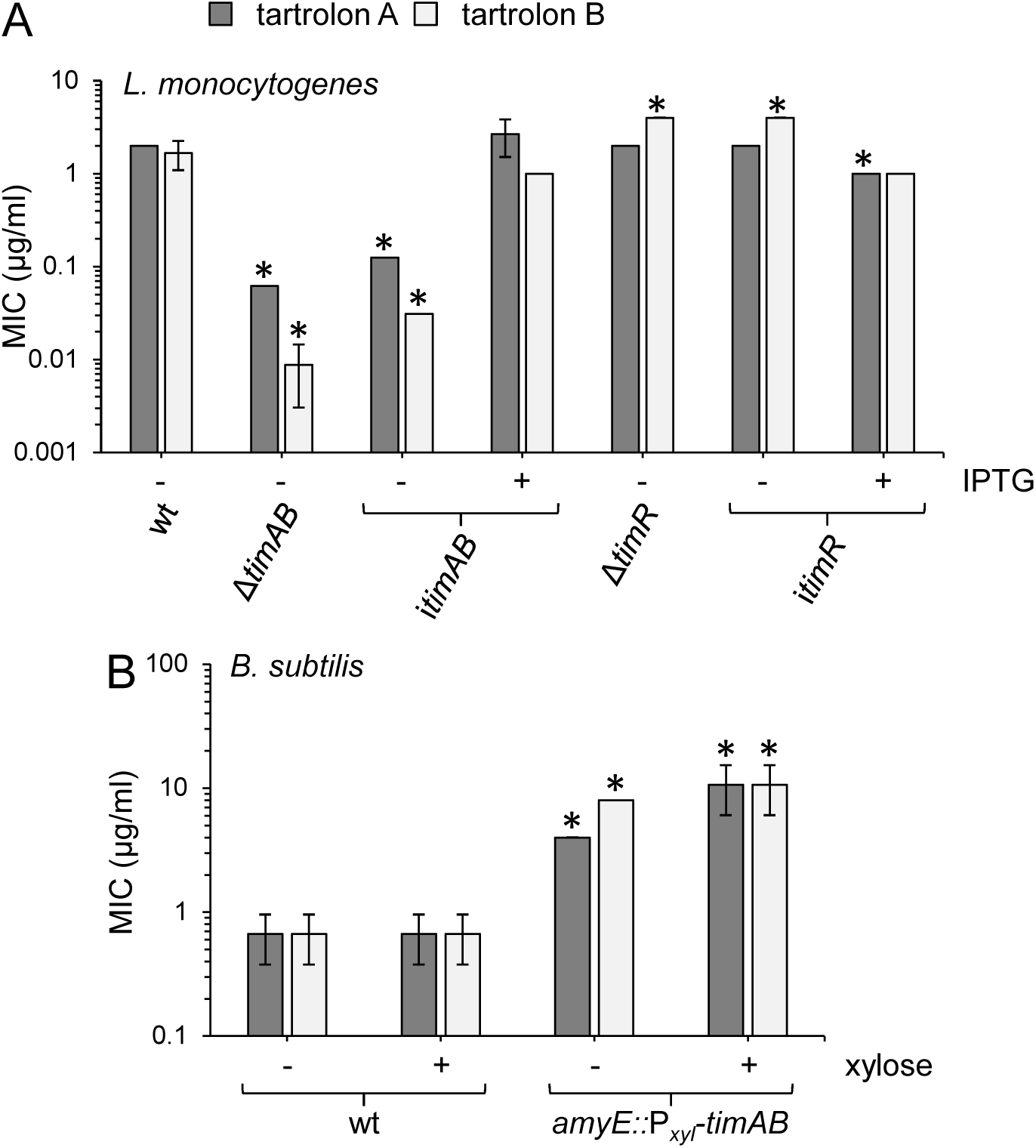
**Contribution of *timABR* genes to tartrolon resistance.** (A) Effect of *timAB* genes on resistance against tartrolon A and B. Minimal inhibitory concentrations (MICs) of *L. monocytogenes* strains EGD-e (wt), LMTE34 (Δ*timAB*), LMTE37 (Δ*timR*), LMTE51 (i*timAB*) and LMTE52 (i*timR*) grown in BHI broth ± 1 mM IPTG were determined in broth dilution assays. (B) Effect of *timAB* expression on tartrolon resistance of *B. subtilis*. MICs of tartrolon A and B were determined for *B. subtilis* strains 168 (wt) and BSTE1 (*amyE::*P*_xyl_-timAB*) grown in LB broth ± 0.5% xylose. Average values and standard deviations were calculated from three independent repetitions. Asterisks mark statistically significant differences compared to wild type (*P*<0.05, *t*-test with Bonferroni-Holm correction).

Next, we asked whether the *timAB* genes would also be sufficient to mediate protection against tartrolon antibiotics. To address this, the *L. monocytogenes timAB* genes were heterologously expressed in *B. subtilis* from a xylose inducible promoter. The MICs of tartrolon A and B were 0.7±0.3 µg/ml for the *B. subtilis* wild type strain 168 in each case. In contrast, strain BSTE1 (*amyE::*P*_xyl_-timAB*) had approximately ten-fold higher MICs for tartrolon A (4±0 µg/ml) and tartrolon B (8±0 µg/ml) and the MICs for both compounds increased even to 10.7±4.4 µg/ml in the presence of xylose (Fig. 4B). In contrast to this, resistance of *B. subtilis* to boromycin was not affected by transplantation and expression of the *timAB* genes (data not shown). This shows that the *timAB* genes encode a novel transporter required, sufficient and specific for resistance against tartrolons.

### TimR binding to the P*_timABR_* promoter is sensitive to tartrolon B

Deletion of *timR* resulted in a four-fold increase of P*_timABR_* promoter activity (Fig. 3C) and also increased tartrolon B resistance (Fig. 4A), which suggested that TimR could repress transcription of its own operon. To further test this hypothesis, TimR was purified as a Strep-tagged protein (Fig. 5A) and binding of TimR to the P*_timABR_* promoter fragment was analysed in an electrophoretic mobility shift assay (EMSA). As can be seen in Fig. 5B, addition of TimR retarded the P*_timABR_* promoter fragment in the gel and two complexes appeared instead, out of which the slower migrating species remained when the TimR concentration was further increased. Most likely, TimR is dimeric as known for other TetR-type transcriptional repressors (41) and increasing concentrations favor TimR dimerization. Intriguingly, no such complex formation was observed when the same experiment was repeated in the presence of tartrolon B, indicating that tartrolon B prevents binding of TimR to the P*_timABR_* promoter fragment (Fig. 5B). In contrast, promoter binding of TimR was neither prevented by tartrolon A or boromycin (Fig. S2), suggesting that TimR is sensitive to the presence of the centrally bound boron atom and to the tartrolon-specific macrocycle.

**Figure 5:**
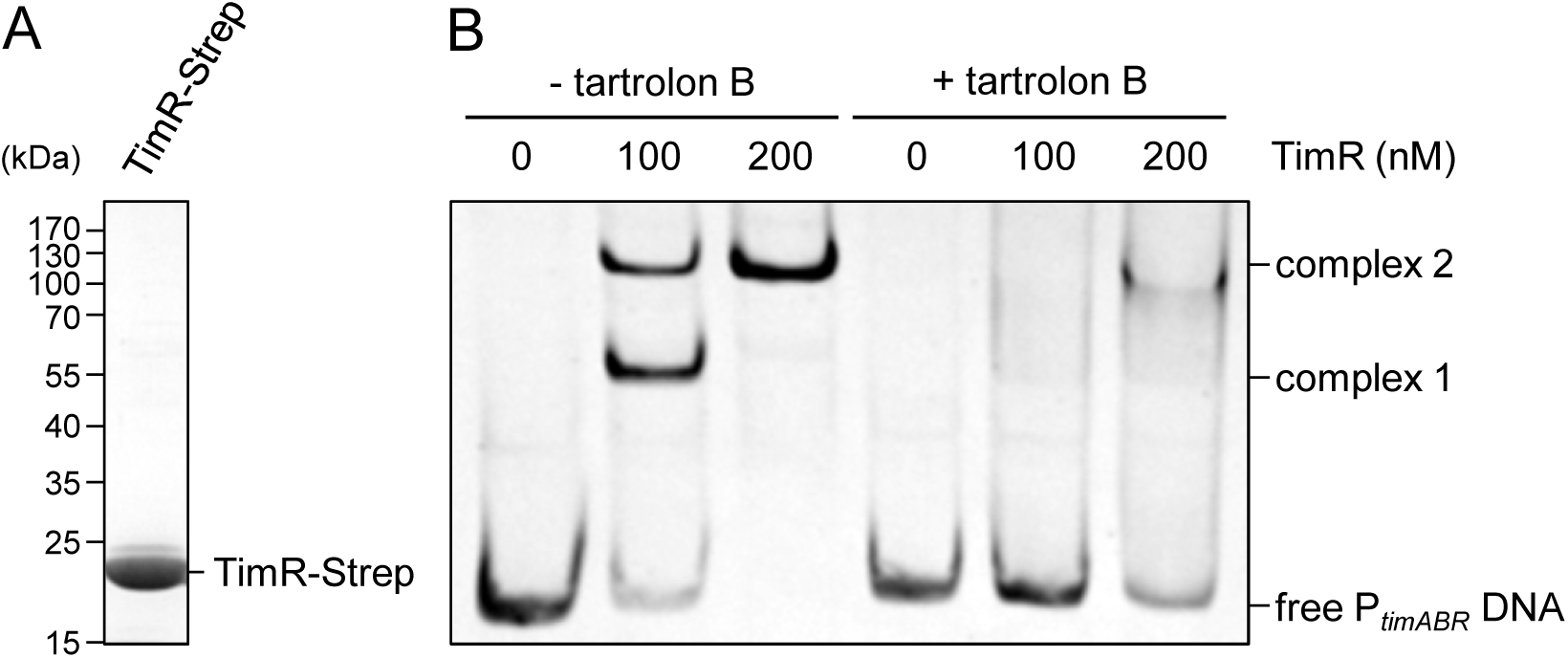
Effect of tartrolon B on TimR binding to the P*_timABR_* promoter (A) SDS-PAGE gel after coomassie staining showing purity of TimR-Strep. (B) Electrophoretic mobility shift assay showing the interaction of TimR-Strep to a P*_timABR_* promoter fragment in the absence and the presence of 1.25 µg/ml tartrolon B. The positions of unbound DNA and the two TimR-promoter nucleoprotein complexes are indicated.

### Evidence for the mode of action of tartrolon B as potassium ionophore

Boromycin acts as a potassium ionophore, causing leakage of potassium ions out of the cell (42, 43). This explains boromycin toxicity, as high intracellular potassium concentrations are critical for pH homeostasis in the cytosol, maintenance of turgor and membrane potential as well as for ribosome activity (44, 45). We wondered, whether the structurally related tartrolon B would act in a similar way and for this we asked whether external potassium supply would suppress efficacy of tartrolon B. As can be seen in Fig. 6, a 200-fold increase of the MIC of boromycin was observed, when 250 mM potassium chloride was added to the growth medium. This is in good agreement with boromycin acting as a potassium ionophore and is consistent with other reports (42, 43). Importantly, a similar effect was also observed for tartrolon B, whose MIC increased 32-fold in the presence of potassium chloride. In contrast, addition of 250 mM sodium chloride did not result in a similar effect, and rather yielded a slight sensitization of *L. monocytogenes* against both antibiotics. This shows that externally supplied potassium ions specifically suppress the toxicity of boromycin and tartrolon B against *L. monocytogenes* and this would be in good agreement with a potassium ionophore mode of action.

**Figure 6:**
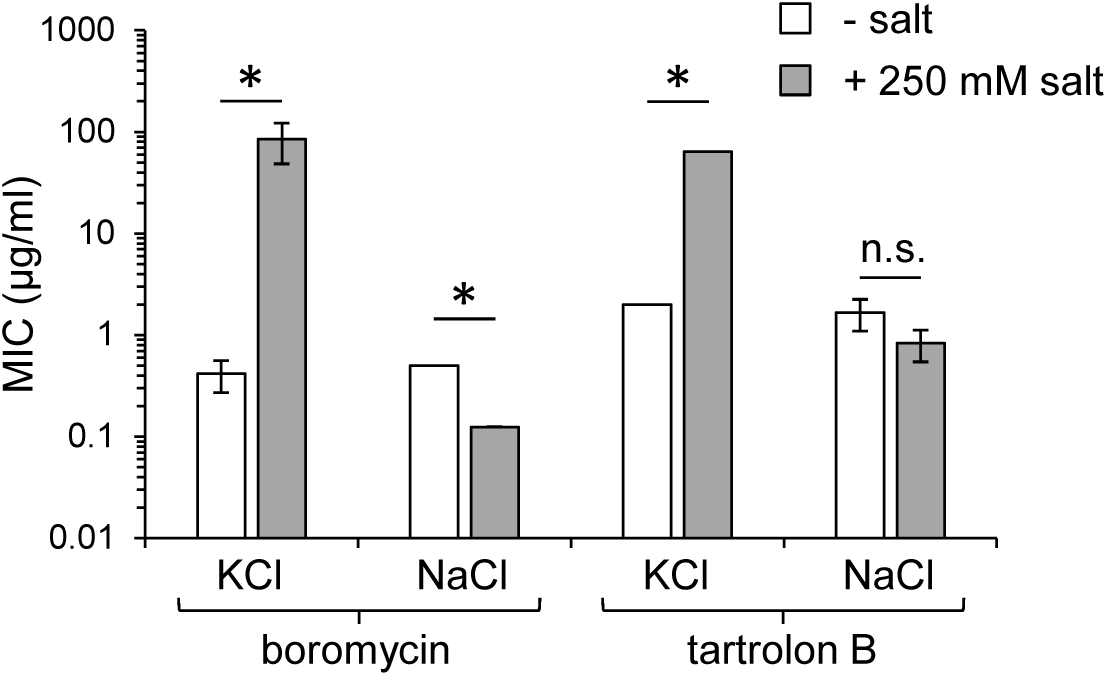
**Tartrolon B action is potassium-sensitive.** MICs of tartrolon B for *L. monocytogenes* EGD-e in the absence and presence of 250 mM KCl. Boromycin (positive control) and NaCl (negative control) were included for comparison. MICs were determined three times and average values with standard deviations are shown. Asterisks mark statistically significant differences (*t*-test, *P*<0.05).

### Tartrolon B resistance of other *Listeria* species

The *timABR* genes are part of the *L. monocytogenes* core genome (46) and they are present in all closed *L. monocytogenes* genomes currently available (data not shown). Likewise, *timABR* homologues are found in all other *Listeria sensu stricto* species (*L. innocua*, *L. ivanovii*, *L. marthii*, *L. seeligeri* and *L. welshimeri*) and in the *Listeria* species belonging to the *Paenilisteria* clade (47), while the *timABR* gene cluster was found to be absent from the genomes of *Listeria* species belonging to the *Mesolisteria* and *Murraya* groups (Fig. S3). Moreover, the *timR* gene of *Listeria cornellensis* FSL F6-0969 (assembly accession: GCA_000525855.1) and the *timB* gene of *Listeria riparia* FSL S10-1204 are truncated (Fig. S3) (48), suggesting that selection for loss or increase of tartrolon resistance is still ongoing or might have occurred parallel to speciation.

We measured tartrolon B resistance of selected *Listeria sensu stricto* (*timABR* positive), *Mesolisteria* (*timABR* negative) and *Murraya* species (*timABR* negative) and found that tartrolon B resistance levels were comparable to *L. monocytogenes* among the three other tested *Listeria sensu stricto species* (Tab. 3) that all contained *timABR* (Fig. S3). Unexpectedly, tartrolon B resistance of *Mesolisteria* species was found to be similar (*L. floridensis*) or even higher (*L. aquatica* and *L. fleischmannii*) as in *L. monocytogenes* (Tab. 3), even though *timABR* was absent (Fig. S3). In contrast, *L. grayi*, representing the *Murraya* clade, was as sensitive to tartrolon B as the *L. monocytogenes* Δ*timAB* mutant (Tab.3, Fig. 4A). The absence of *timABR* from the *L. grayi* genome would explain this sensitivity, but *timABR*-independent tarrtolon B resistance mechanisms must have evolved in *Mesolisteria*.

**Table 3:**
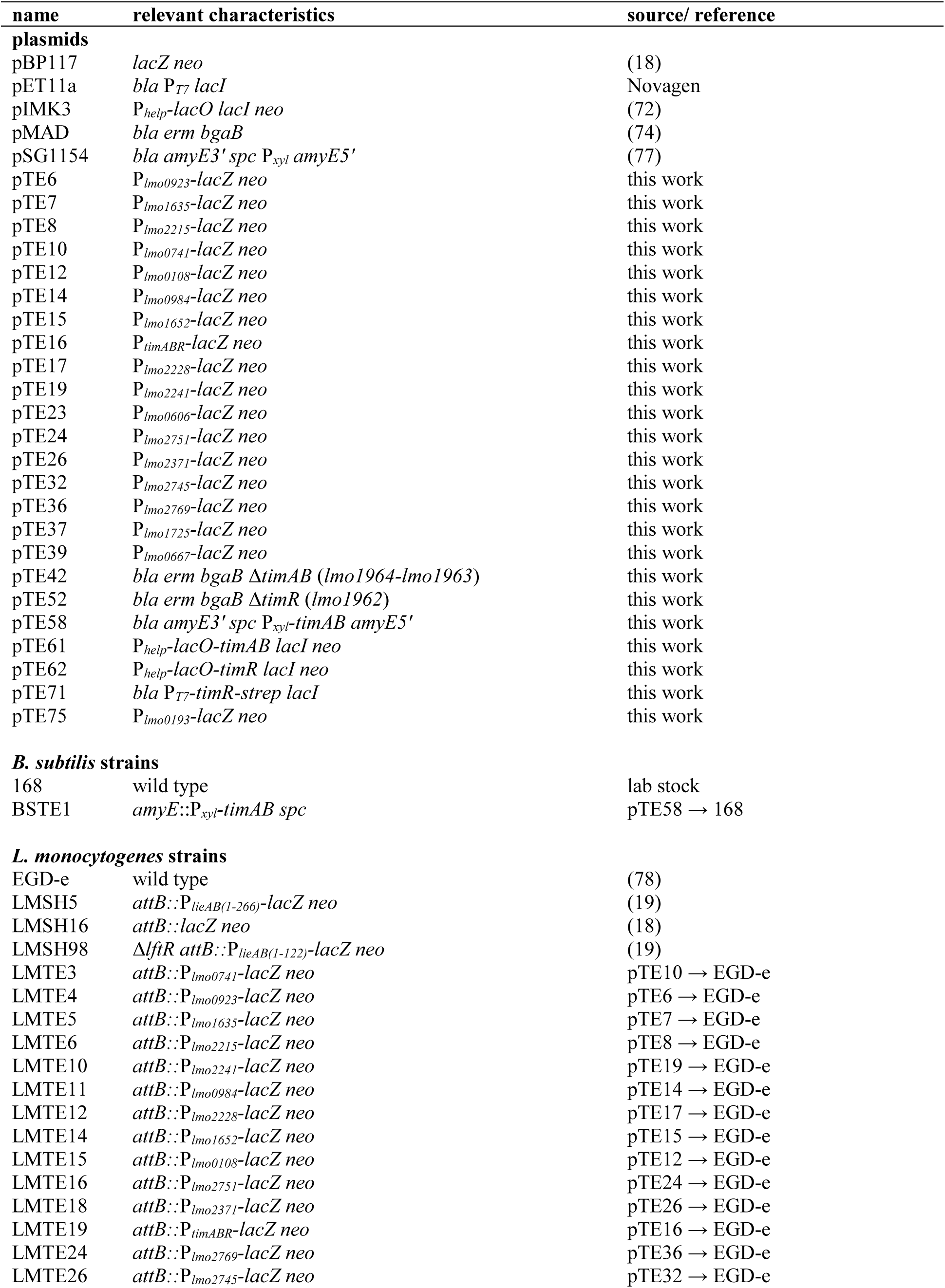
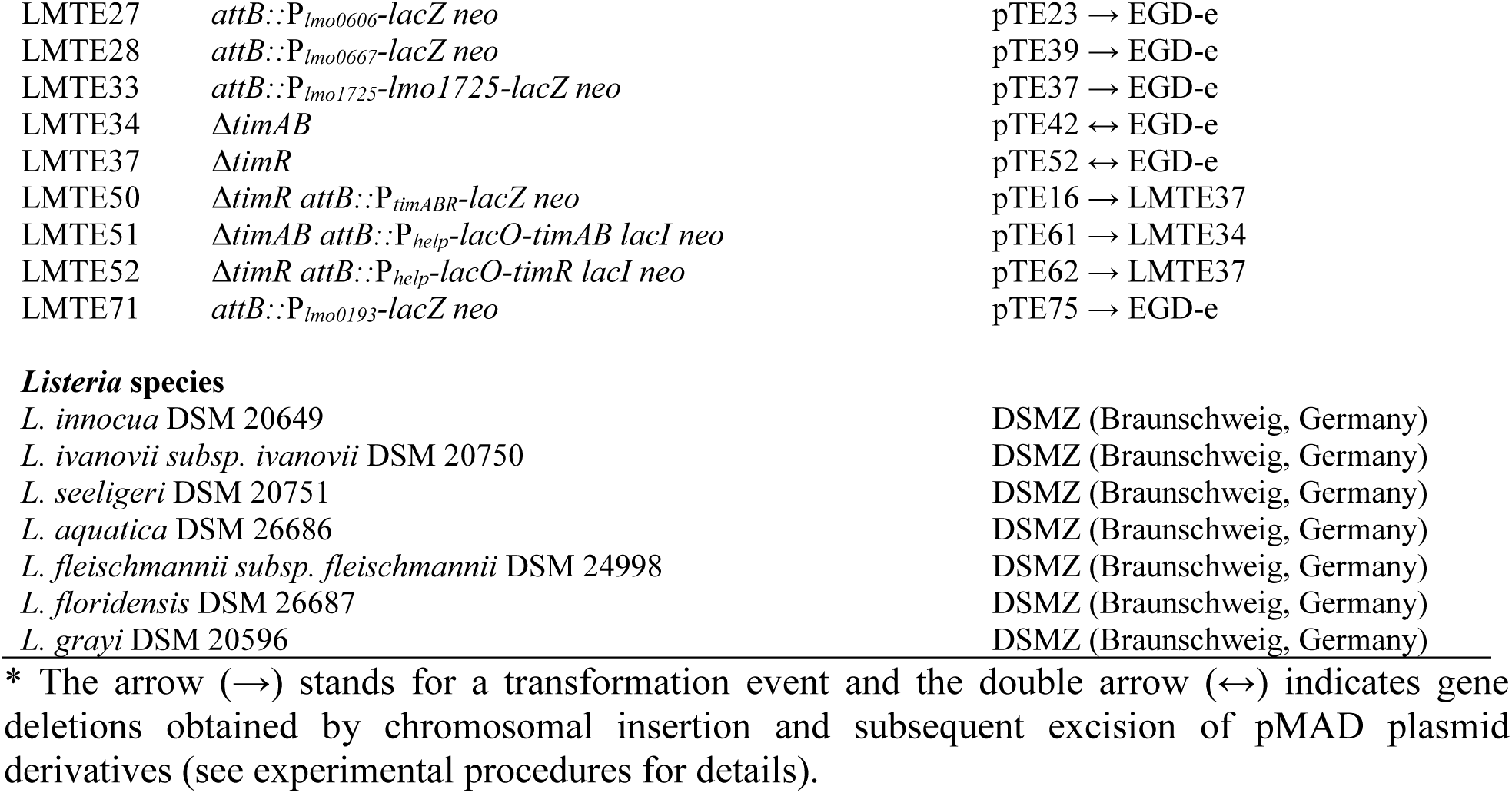
Plasmids and strains used in this study.

## Discussion

As a ubiquitously occurring environmental bacterium, *L. monocytogenes* has developed manifold strategies to survive the conditions prevailing in its multifaceted habitats. The stress response mechanisms of *L. monocytogenes* facing those harmful conditions that are most obviously occurring during life in the environment such as heat/cold, high osmolarity or exposure to ultraviolet light are generally well understood (5, 49, 50). The response to conditions specific to the passage through the gastrointestinal tract including the extreme acidity in the stomach or the presence of bile salts in the gut has also been investigated (51, 52). In contrast, studies on how *L. monocytogenes* interacts with or outcompetes other species of the gut microbiota by production of bacteriocins such as listeriolysin S or Lmo2776 began to move into the focus of research just recently (53-55), even though research on listerial resistance against such bacteriocins, particularly nisin, which is used as a food preservative, has a long tradition (56). Despite intensive investigations of the confrontation of *L. monocytogenes* with this wide spectrum of noxious conditions/agents, there is only limited knowledge on the strategies used by *L. monocytogenes* to withstand antibiotics produced by rivaling species in habitats outside the human host.

One example for systems providing resistance against antibiotics produced by natural competitors is the LftRS/LieAB system enabling *L. monocytogenes* to sense and to detoxify aurantimycin A (18, 19), which is produced by *S. aurantiacus*, another inhabitant of the soil (24). With the TimABR system, we here add another example to the list of *L. monocytogenes* systems providing resistance against an antibiotic synthesized by a soil-dwelling microorganism co-residing in the same habitat. Similar to the LftRS/LieAB system, this system is composed of an ABC transporter for compound export and a transcriptional regulator for compound sensing. While direct sensing of aurantimycin by its cognate transcriptional regulator LftR has never been shown (18, 19), a direct interaction of tartrolon B with TimR, the transcriptional regulator of the TimABR system, must be concluded from the observation that tartrolon B addition prevents promoter binding of TimR. Remarkably, this interaction was specific to the borate-containing tartrolon B and could not be observed with the identical but borate-free macrodiolide ring present in tartrolon A. The conformation of the macrocycle in tartrolon B and thus its ability to bind to TimR might be altered after borate esterification. Alternatively, TimR could make direct contacts with the borate ester part of the tartrolon B molecule through electrostatic interactions or could form hydrogen bonds with the borate oxygens as observed in *Vibrio harveyi* LuxP, the sensor of the borate diester containing quorum sensing autoinducer-2 (57). Interestingly, even covalent bonds between boron atoms of organoboron compounds and amino acid side chains of proteins have been reported (58, 59) and could potentially be involved in tartrolon B recognition by TimR. Despite this remarkable specificity of TimR for tartrolon B, tartrolon detoxification by TimAB is not sensitive to the presence of the borate ester, as judged from the *timAB* transplantation experiment into *B. subtilis*. That the effect of a *timAB* deletion on the MIC of tartrolon A is less pronounced than that for tartrolon B (Fig. 4A), could also be explained by the less efficient induction of the P*_timABR_* promoter by tartrolon A compared to tartrolon B.

Tartrolons were identified in the soil-dwelling myxobacterium *S. cellulosum* (38, 39), from a *Streptomyces* isolate found in a marine sediment (60) and from *Teredinibacter turnerae*, a γ- proteobacterium isolated from the gills of marine shipworms (61). *L. monocytogenes* had frequently been isolated from soil samples of various origin (62), its isolation from seawater and estuarine water samples was reported at least occasionally (63-65) and very often the pathogen is found on seafood products (66, 67). *L. monocytogenes* tolerates high salinity (68, 69), and therefore isolation from marine environments seems plausible. The strong conservation of the *timABR* locus in selected *Listeria* lineages could reflect a tight ecological interaction between tartrolon-producing microorganisms present in the soil and in marine environments and such *timABR*-positive *Listeria* species.

The *timABR* cassette is only one out of the 18 ABC transporter operons with which we started this project. The inducing agents and/or the compounds exported by the remaining ones are still unknown. We assume that the specific compilation of these still to be characterized ABC transporters represents a genetic signature that reflects the ecological interactions that *L. monocytogenes* experiences with its most important antibiotic-producing competitors in nature. Understanding these interactions in combination with knowledge on the prevalence of the compound producers in nature might help to better specify the environmental reservoir of *L. monocytogenes*.

## Materials and Methods

### Bacterial strains and growth conditions

All strains and plasmids used in this study are listed in Tab. 3. Non-*L. monocytogenes Listeria* strains were received from the German Collection of Microorganisms and Cell Cultures (DSMZ, Braunschweig). *Listeria* strains were grown in BHI broth or on BHI agar plates at 37°C. Strains of *B. subtilis* were cultivated in LB broth or on LB agar plates. Erythromycin (5 µg ml^-1^), kanamycin (50 µg ml^-1^), spectinomycin (50 µg ml^-1^), X-Gal (100 µg ml^-1^) or IPTG (1 mM) were added as indicated where required. *Escherichia coli* TOP10 was used as the standard cloning host (70). Tartrolon A and B were obtained from the DZIF natural compound selection, boromycin was purchased from Hello Bio Ltd. (Ireland).

### General methods, manipulation of DNA and oligonucleotide primers

Standard methods were used for transformation of *E. coli* and for isolation of plasmid DNA (70). Transformation of *L. monocytogenes* and *B. subtilis* was carried out as described by others (71, 72). Restriction and ligation of DNA was performed following the manufactureŕs instructions. All primer sequences are listed in Tab. 4.

**Table 4:**
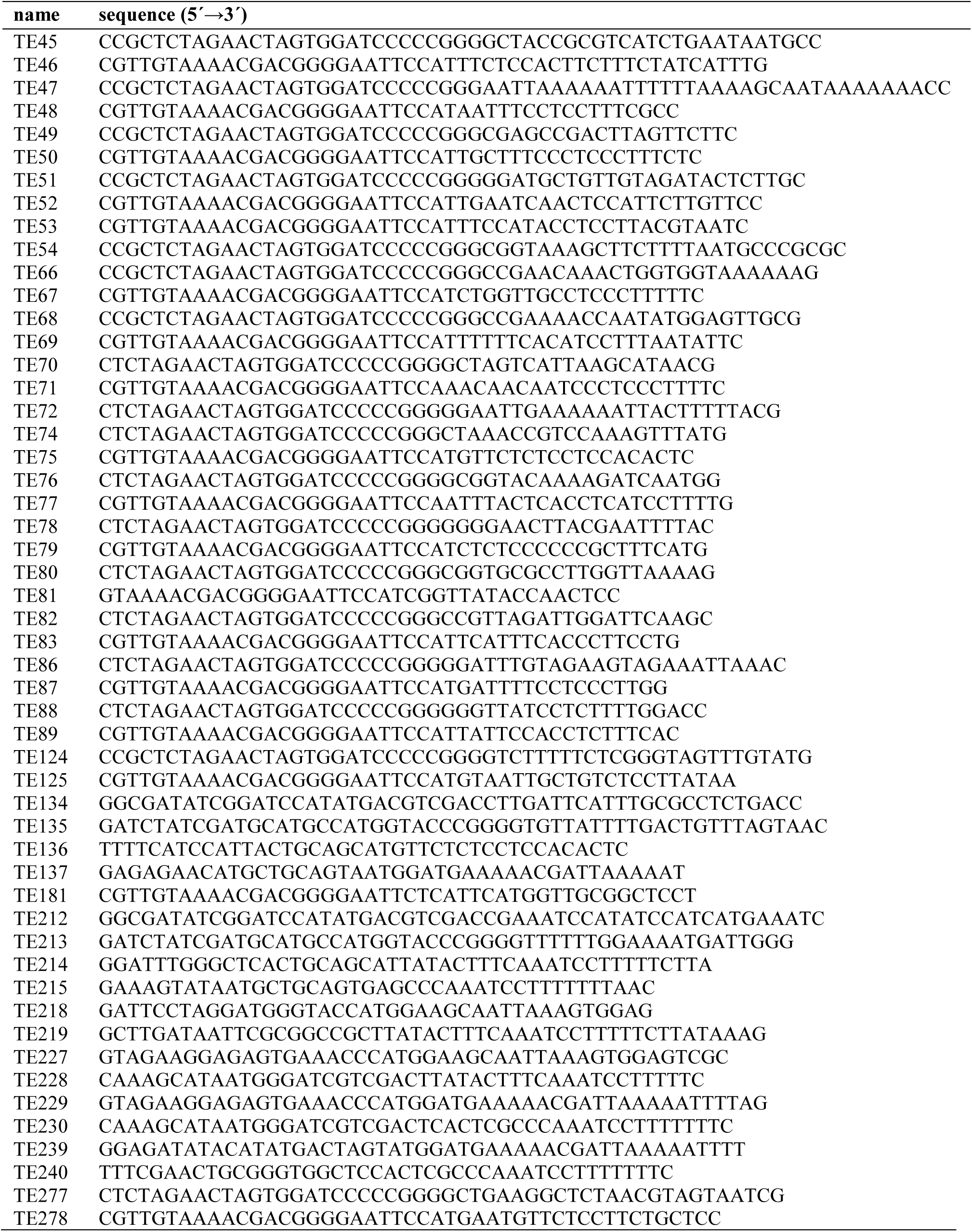
Primers used in this study.

### Construction of bacterial plasmids and strains

Plasmids carrying *lacZ* fusions to the promoters of ABC transporter operons were constructed by amplification of the promotor sequences using primers TE45/TE46 (P*_lmo0606_*), TE47/TE48 (P*_lmo0741_*), TE49/TE50 (P*_lmo0923_*), TE51/TE52 (P*_lmo1635_*), TE53/TE54 (P*_lmo2215_*), TE66/TE67 (P*_lmo0984_*), TE68/TE69 (P*_lmo0108_*), TE70/TE71 (P*_lmo1652_*), TE72/TE181 (P*_lmo1725_*), TE74/TE75 (P*_timABR_*), TE76/TE77 (P*_lmo2228_*), TE78/TE79 (P*_lmo2241_*), TE80/TE81 (P*_lmo2371_*), TE82/TE83 (P*_lmo2745_*), TE86/TE87 (P*_lmo2751_*), TE88/TE89 (P*_lmo2769_*), TE124/TE125 (P*_lmo0667_*) or TE277/TE278 (P*_lmo0193_*).

The resulting fragments were inserted into pBP117 using restriction free (RF) cloning (73).

For construction of plasmid pTE42, allowing deletion of *timAB*, fragments up-and downstream to *timAB* were amplified using oligonucleotides TE135/TE136 and TE137/TE134, respectively. Both fragments were then fused together by splicing by overlapping extension (SOE)-PCR and introduced into pMAD by RF cloning. Plasmid pTE52, designed for deletion of *timR*, was obtained by amplification of *timR* up-and downstream fragments using the primer pairs TE213/TE214 and TE215/TE212, respectively. These fragments were spliced together by SOE-PCR and the resulting fragment was introduced into pMAD by RF cloning.

Plasmid pTE61 was constructed for inducible *timAB* expression. It was obtained by amplification of *timAB* using the oligonucleotides TE227/TE228 and cloning of the resulting fragment into pIMK3 using NcoI/SalI.

Plasmid pTE62 was constructed for IPTG-dependent *timR* expression. To this end, the *timR* gene was amplified using TE229/TE230 as the primers and inserted into pIMK3 using NcoI/SalI. Plasmid pTE71 was generated for purification of strep-tagged TimR. For this, the *timR* gene was amplified with oligonucleotides TE239/TE240 and integrated into pET11a through RF cloning.

Plasmid pTE58 was constructed for heterologous *timAB* expression in *B. subtilis*. To this end, the *timAB* genes were amplified using the primer pair TE218/TE219 and the resulting fragment was cloned into pSG1154 using RF cloning.

Derivatives of pIMK3 and pBP117 plasmids were introduced into *L. monocytogenes* strains by electroporation and transformants were selected on BHI agar plates containing kanamycin at 37°C. Plasmid integration at the tRNA^Arg^ *attB* site was confirmed by PCR. Likewise, plasmid derivatives of pMAD were introduced into *L. monocytogenes*, but transformants were selected on BHI agar plates containing X-Gal and erythromycin at 30°C. The plasmid integration-excision protocol described by Arnaud *et al.* (74) was then used for gene deletions. All gene deletions were confirmed by PCR. *B. subtilis* was transformed with plasmid pTE58 and transformants were selected on LB agar plates containing spectinomycin (50 µg/ml). Integration of the plasmid into the *amyE* locus was confirmed by absence of amylase activity on starch containing agar plates.

### Natural compound screen

Natural compounds were compiled by the German Centre for Infection Research (DZIF) infrastructure and provided as “Natural Compound Library” through the Helmholtz Institute for Pharmaceutical Research Saarland (HIPS) as a ready-to-screen library (37). The compilation includes 681 purified secondary metabolites from myxobacteria (253), fungi (88), and streptomycetes (340), collected in natural product screening programs at HIPS, the Helmholtz Centre for Infection Research (HZI), and the University of Tübingen, respectively. Library compounds were provided as 1 mM stock solutions in DMSO in conical 96-well plates in a randomized order and encrypted by a barcode system for non-biased screening. For the screen, *L. monocytogenes* reporter strains were grown over night in BHI broth at 37°C and diluted 1:2000 in molten BHI agar containing 50 µg/ml X-Gal, which was cooled down to ∼50°C prior to mixing and plate pouring. Compounds were applied on top of the agar plates (1 µl each) at a concentration of 0.033 mM (in DMSO) and plates were incubated over night at 37°C.

### Compound quality control

Compounds included in the natural product library are routinely checked by HPLC-MS and NMR. Here, we additionally analyzed the batches of tartrolons A and B that were used for extended biological assays. Measurements were performed with a Dionex Ultimate 3000 RSLC system (Thermo) using a BEH C18, 100 × 2.1 mm, 1.7 µm dp column (Waters). Separation of 1 µl sample was achieved by a linear gradient from (A) H_2_O + 0.1% FA to (B) ACN + 0.1% FA at a flow rate of 600 µl/min and 45 °C. The gradient was initiated by a 0.5 min isocratic step at 5% B, followed by an increase to 95% B in 18 min to end with a 2 min step at 95% B before reequilibration with initial conditions. UV spectra were recorded by a diode array detector (DAD) in the range from 200 to 600 nm. The LC flow was split to 75 µl/min before entering the timsTOF fleX mass spectrometer (Bruker Daltonics). The split was set up with fused silica capillaries of 75 and 100 µm I.D. and a low dead volume tee junction (Upchurch). The timsTOF fleX was operated in positive ESI mode, with 1.0 bar nebulizer pressure, 5.0 l/min dry gas, 200°C dry heater, 4000 V capillary voltage, 500 V end plate offset, 600 Vpp funnel 1 RF, 400 Vpp funnel 2 RF, 80 V deflection delta, 5 eV ion energy, 10 eV collision energy, 1500 Vpp collision RF, 10 µs pre-pulse storage, 100 µs transfer time. TIMS (trapped ion mobility spectrometry) delta values were set to -20 V (delta 1), -120 V (delta 2), 80 V (delta 3), 100 V (delta 4), 0 V (delta 5), and 100 V (delta 6). The 1/k_0_ (inverse reduced ion mobility) range was set from 0.55 Vs/cm^2^ to 1.9 Vs/cm^2^, the mass range was *m/z* 100-2000. Ion charge control (ICC) was enabled and set to 7.5 million counts. The samples were analyzed with TIMS ramp times of 100 ms. The analysis accumulation and ramp time was set at 100 ms with a spectra rate of 9.43 Hz and a total cycle of 0.32 sec was also selected resulting in one full TIMS-MS scan. TIMS dimension was calibrated linearly using 4 selected ions from ESI Low Concentration Tuning Mix (Agilent Technologies) [*m/z*, 1/k_0_: (301.998139, 0.6678 Vs/cm^2^), (601.979077, 0.8782 Vs/cm^2^)] in negative mode and [*m/z*, 1/k_0_: (322.048121, 0.7363 Vs/cm^2^), (622.028960, 0.9915 Vs/cm^2^)] in positive mode. The mobility-mass correlation for calibration was taken from the CCS compendium (75).

### β-galactosidase assay

For measurement of the activity of promoters fused to *lacZ*, an overnight culture of each strain was diluted 1:100 in 5 ml BHI broth and compounds added where needed. Strains were grown at 37°C and 250 rpm until an OD_600_ of 0.6 – 0.8 and pelleted by centrifugation (11000 x g, 2 min). The pellet was washed once in 600 µl ddH_2_O and resuspended in 1200 µl PBS (137 mM NaCl, 2.7 mM KCl, 10 mM phosphate buffer, pH 7.4) buffer containing 0.15% β-mercaptoethanol. After sonification and centrifugation (11000 x g, 5 min), 1000 µl of the supernatant was incubated at 30°C for 10 minutes. 200 µl 4 mg/ml ONPG in 1x PBS was added and the reaction was incubated at 30°C for another 10 minutes, before being stopped by addition of 500 µl 1 M Na_2_CO_3_. Absorption was measured at 420 nm against PBS buffer incubated with ONPG as blank value. Protein concentration was determined by mixing 50 µl of the supernatant and 950 µl of a 1 x Roti®Nanoquant (Carl Roth, Karlsruhe, Germany) solution and measurement of the absorption at 595 nm against 50 µl 1x PBS as blank value. Afterwards, the promoter activity in Miller units (MU) was calculated.

### Determination of minimal inhibitory concentrations

Minimal inhibitory concentrations (MIC) were determined in a 96-well plates in a total volume of 200 µl. 200 µl BHI containing a defined dilution series of the antibiotic of interest were inoculated with overnight cultures at an initial OD_600_ of 0.05. The microtiter plates were incubated in a plate reader with intermittent shaking over night at 37°C and growth was recorded for 20 h. The MIC was defined as the lowest concentration of the antibiotic at which no growth could be observed.

### Protein purification

Overexpression of *timR-strep* was performed in *E. coli* BL21. To this end, 500 ml LB broth containing 100 µg/ml ampicillin were inoculated with *E. coli* BL21 cells carrying the corresponding vector construct to an initial OD_600_ of 0.1. Cells were grown at 37 °C and 250 rpm and 1 mM isopropyl-β-d-thiogalactopyranoside (IPTG) was added when an OD_600_ of 0.6 – 0.8 was reached. After three more hours, cells were pelleted by centrifugation (6,000 × *g*, 5 min, 4°C) and the pellet was washed once with 25 ml buffer W (100 mM Tris-HCl pH 8.0; 150 mM NaCl). Cells were resuspended in 40 ml buffer W and lysed using an Emulsiflex homogenizer (Avestin, Germany). Cell debris was removed by centrifugation (6,000 × *g*, 5 min, 4°C), and the resulting supernatant was filtered through a Minisart filter with a pore size of 0.45 μm (Sartorius). The strep tagged protein was purified using affinity chromatography and Strep-Tactin Sepharose (IBA Lifesciences, Germany) according to the manufacturer’s instructions. Fractions containing purified proteins were pooled, aliquoted, and stored at –20°C. Samples purity was analyzed by standard sodium dodecyl sulfate polyacrylamide gel electrophoresis (SDS-PAGE) followed by Coomassie staining.

### Electrophoretic mobility shift assay (EMSA)

To test the interaction of TimR with the P*_timABR_* promoter, a 450 bp long sequence upstream of *timA* was synthesized by PCR from EGD-e genomic DNA using primers TE74 and TE75 and purified using the NucleoSpin® Gel and PCR Clean-up Kit (Macherey-Nagel, Düren, Germany). For the EMSA experiment, 32 nM of this purified promoter fragment was mixed with 5 µl of EMSA-buffer (120 mM HEPES, 300 mM KCl, 30 mM MgCl_2_, 0.3 mg/ml bovine serum albumin [BSA], 30% glycerol, 0.3 mM EDTA, pH 8.0) and incubated with varying amounts of TimR-Strep, tartrolon A, tartrolon B and boromycin for 5 minutes at room temperature. The reaction was loaded onto an acrylamide gel (10 % (w/v) acrylamide with 0.6 x Tris-borate EDTA) and run at 120 V for 75 minutes. Afterwards, the gel was stained for 5 minutes using ethidium bromide and photographed using an UV transilluminator.

## Acknowledgements

This work was supported by a grant of the DFG (HA6830/2-1) to S. H. The authors are grateful to all group members for fruitful discussions. We would also like to thank Kerstin Schober and Viktoria George for excellent technical assistance with natural products supply and Sabine Backes, Dr. Jake Haeckl and Dr. Susanne Kirsch-Dahmen for support of analytics.

## References

1. Truong HN, Garmyn D, Gal L, Fournier C, Sevellec Y, Jeandroz S, Piveteau P. 2021. Plants as a realized niche for Listeria monocytogenes. Microbiologyopen 10:e1255.

2. Freitag NE, Port GC, Miner MD. 2009. *Listeria monocytogenes* - from saprophyte to intracellular pathogen. Nat Rev Microbiol 7:623–8.

3. Esteban JI, Oporto B, Aduriz G, Juste RA, Hurtado A. 2009. Faecal shedding and strain diversity of *Listeria monocytogenes* in healthy ruminants and swine in Northern Spain. BMC Vet Res 5:2.

4. Schoder D, Guldimann C, Martlbauer E. 2022. Asymptomatic Carriage of *Listeria monocytogenes* by Animals and Humans and Its Impact on the Food Chain. Foods 11.

5. Quereda JJ, Moron-Garcia A, Palacios-Gorba C, Dessaux C, Garcia-Del Portillo F, Pucciarelli MG, Ortega AD. 2021. Pathogenicity and virulence of *Listeria monocytogenes*: A trip from environmental to medical microbiology. Virulence 12:2509–2545.

6. Vazquez-Boland JA, Kuhn M, Berche P, Chakraborty T, Dominguez-Bernal G, Goebel W, Gonzalez-Zorn B, Wehland J, Kreft J. 2001. *Listeria* pathogenesis and molecular virulence determinants. Clin Microbiol Rev 14:584–640.

7. Charlier C, Perrodeau E, Leclercq A, Cazenave B, Pilmis B, Henry B, Lopes A, Maury MM, Moura A, Goffinet F, Dieye HB, Thouvenot P, Ungeheuer MN, Tourdjman M, Goulet V, de Valk H, Lortholary O, Ravaud P, Lecuit M, group Ms. 2017. Clinical features and prognostic factors of listeriosis: the MONALISA national prospective cohort study. Lancet Infect Dis 17:510–519.

8. Wilking H, Lachmann R, Holzer A, Halbedel S, Flieger A, Stark K. 2021. Ongoing High Incidence and Case-Fatality Rates for Invasive Listeriosis, Germany, 2010-2019. Emerg Infect Dis 27:2485–2488.

9. Tiensuu T, Guerreiro DN, Oliveira AH, O’Byrne C, Johansson J. 2019. Flick of a switch: regulatory mechanisms allowing *Listeria monocytogenes* to transition from a saprophyte to a killer. Microbiology 165:819–833.

10. Lubelski J, Konings WN, Driessen AJ. 2007. Distribution and physiology of ABC-type transporters contributing to multidrug resistance in bacteria. Microbiol Mol Biol Rev 71:463–76.

11. Mata MT, Baquero F, Perez-Diaz JC. 2000. A multidrug efflux transporter in *Listeria monocytogenes*. FEMS Microbiol Lett 187:185–8.

12. Godreuil S, Galimand M, Gerbaud G, Jacquet C, Courvalin P. 2003. Efflux pump Lde is associated with fluoroquinolone resistance in *Listeria monocytogenes*. Antimicrob Agents Chemother 47:704–8.

13. Müller A, Rychli K, Zaiser A, Wieser C, Wagner M, Schmitz-Esser S. 2014. The *Listeria monocytogenes* transposon Tn6188 provides increased tolerance to various quaternary ammonium compounds and ethidium bromide. FEMS Microbiol Lett 361:166–73.

14. Young J, Holland IB. 1999. ABC transporters: bacterial exporters-revisited five years on. Biochim Biophys Acta 1461:177–200.

15. Wendlandt S, Shen J, Kadlec K, Wang Y, Li B, Zhang WJ, Fessler AT, Wu C, Schwarz S. 2015. Multidrug resistance genes in staphylococci from animals that confer resistance to critically and highly important antimicrobial agents in human medicine. Trends Microbiol 23:44–54.

16. Sleator RD, Wemekamp-Kamphuis HH, Gahan CG, Abee T, Hill C. 2005. A PrfA-regulated bile exclusion system (BilE) is a novel virulence factor in *Listeria monocytogenes*. Mol Microbiol 55:1183–95.

17. Collins B, Curtis N, Cotter PD, Hill C, Ross RP. 2010. The ABC transporter AnrAB contributes to the innate resistance of *Listeria monocytogenes* to nisin, bacitracin, and various beta-lactam antibiotics. Antimicrob Agents Chemother 54:4416–23.

18. Hauf S, Herrmann J, Miethke M, Gibhardt J, Commichau FM, Muller R, Fuchs S, Halbedel S. 2019. Aurantimycin resistance genes contribute to survival of *Listeria monocytogenes* during life in the environment. Mol Microbiol 111:1009–1024.

19. Hauf S, Engelgeh T, Halbedel S. 2021. Elements in the LftR Repressor Operator Interface Contributing to Regulation of Aurantimycin Resistance in *Listeria monocytogenes*. J Bacteriol 203.

20. Jiang X, Geng Y, Ren S, Yu T, Li Y, Liu G, Wang H, Meng H, Shi L. 2019. The VirAB-VirSR-AnrAB Multicomponent System Is Involved in Resistance of *Listeria monocytogenes* EGD-e to Cephalosporins, Bacitracin, Nisin, Benzalkonium Chloride, and Ethidium Bromide. Appl Environ Microbiol 85.

21. Kaval KG, Hahn B, Tusamda N, Albrecht D, Halbedel S. 2015. The PadR-like transcriptional regulator LftR ensures efficient invasion of *Listeria monocytogenes* into human host cells. Front Microbiol 6:772.

22. Lee C, Kim MI, Park J, Hong M. 2019. Structure-based molecular characterization and regulatory mechanism of the LftR transcription factor from *Listeria monocytogenes*: Conformational flexibilities and a ligand-induced regulatory mechanism. PLoS One 14:e0215017.

23. Grigoriev P, Schlegel R, Dornberger K, Gräfe U. 1995. Formation of membrane pores by aurantimycins A and B, new lipopeptide antibiotics from *Streptomyces aurantiacus*. Bioelectrochemistry and Bioenergetics 36:57–59.

24. Gräfe U, Schlegel R, Ritzau M, Ihn W, Dornberger K, Stengel C, Fleck WF, Gutsche W, Härtl A, Paulus EF. 1995. Aurantimycins, new depsipeptide antibiotics from *Streptomyces aurantiacus* IMET 43917. Production, isolation, structure elucidation, and biological activity. J Antibiot (Tokyo) 48:119–25.

25. Aase B, Sundheim G, Langsrud S, Rorvik LM. 2000. Occurrence of and a possible mechanism for resistance to a quaternary ammonium compound in *Listeria monocytogenes*. Int J Food Microbiol 62:57–63.

26. Kopp F, Marahiel MA. 2007. Macrocyclization strategies in polyketide and nonribosomal peptide biosynthesis. Nat Prod Rep 24:735–49.

27. Lubelski J, Rink R, Khusainov R, Moll GN, Kuipers OP. 2008. Biosynthesis, immunity, regulation, mode of action and engineering of the model lantibiotic nisin. Cell Mol Life Sci 65:455–76.

28. Vijayabharathi R, Bruheim P, Andreassen T, Raja DS, Devi PB, Sathyabama S, Priyadarisini VB. 2011. Assessment of resistomycin, as an anticancer compound isolated and characterized from *Streptomyces aurantiacus* AAA5. J Microbiol 49:920–6.

29. Saraylou M, Nadian Ghomsheh H, Enayatizamir N, Rangzan N, St. Clair Senn S. 2021. Some plant growth promoting traits of Streptomyces species isolated from various crop rhizospheres with high root colonization ability of spinach (Spinacia oleracea L.). Applied Ecology and Environmental Research 19:3069–3081.

30. Cavanagh D, Fitzgerald GF, McAuliffe O. 2015. From field to fermentation: the origins of *Lactococcus lactis* and its domestication to the dairy environment. Food Microbiol 47:45–61.

31. Scheldeman P, Herman L, Foster S, Heyndrickx M. 2006. *Bacillus sporothermodurans* and other highly heat-resistant spore formers in milk. J Appl Microbiol 101:542–55.

32. Ren Q, Chen K, Paulsen IT. 2007. TransportDB: a comprehensive database resource for cytoplasmic membrane transport systems and outer membrane channels. Nucleic Acids Res 35:D274–9.

33. Ren Q, Kang KH, Paulsen IT. 2004. TransportDB: a relational database of cellular membrane transport systems. Nucleic Acids Res 32:D284–8.

34. Jain D. 2015. Allosteric control of transcription in GntR family of transcription regulators: A structural overview. IUBMB Life 67:556–63.

35. Housseini BIK, Phan G, Broutin I. 2018. Functional mechanism of the efflux pumps transcription regulators from *Pseudomonas aeruginosa* based on 3D structures. Front Mol Biosci 5:57.

36. Gupta A, Pande A, Sabrin A, Thapa SS, Gioe BW, Grove A. 2019. MarR family transcription factors from *Burkholderia* species: Hidden clues to control of virulence-associated genes. Microbiol Mol Biol Rev 83.

37. DZIF/TTU9. 2023. Compound Resources and Medicinal Chemistry. https://www.dzif.de/en/compound-resources-and-medicinal-chemistry. Accessed

38. Schummer D, Irschik H, Reichenbach H, Höfle G. 1994. Antibiotics from gliding bacteria, LVII. Tartrolons: New boron-containing macrodiolides from *Sorangium cellulosum*. Liebigs Annalen der Chemie 1994:283–289.

39. Irschik H, Schummer D, Gerth K, Hofle G, Reichenbach H. 1995. The tartrolons, new boron-containing antibiotics from a myxobacterium, Sorangium cellulosum. J Antibiot (Tokyo) 48:26–30.

40. Hütter R, Keller-Schien W, Knüsel F, Prelog V, Rodgers jr. GC, Suter P, Vogel G, Voser W, Zähner H. 1967. Stoffwechselprodukte von Mikroorganismen. 57. Mitteilung. Boromycin. Helvetica Chimica Acta 50:1533–1539.

41. Bertram R, Neumann B, Schuster CF. 2021. Status quo of tet regulation in bacteria. Microb Biotechnol doi:10.1111/1751-7915.13926.

42. Moreira W, Aziz DB, Dick T. 2016. Boromycin Kills Mycobacterial Persisters without Detectable Resistance. Front Microbiol 7:199.

43. Pache W, Zähner H. 1969. Metabolic products of microorganisms. 77. Studies on the mechanism of action of boromycin. Arch Mikrobiol 67:156–65.

44. Epstein W. 2003. The roles and regulation of potassium in bacteria. Prog Nucleic Acid Res Mol Biol 75:293–320.

45. Gundlach J, Herzberg C, Hertel D, Thurmer A, Daniel R, Link H, Stülke J. 2017. Adaptation of Bacillus subtilis to Life at Extreme Potassium Limitation. mBio 8.

46. Ruppitsch W, Pietzka A, Prior K, Bletz S, Fernandez HL, Allerberger F, Harmsen D, Mellmann A. 2015. Defining and Evaluating a Core Genome Multilocus Sequence Typing Scheme for Whole-Genome Sequence-Based Typing of *Listeria monocytogenes*. J Clin Microbiol 53:2869–76.

47. Orsi RH, Wiedmann M. 2016. Characteristics and distribution of Listeria spp., including *Listeria* species newly described since 2009. Appl Microbiol Biotechnol 100:5273–87.

48. den Bakker HC, Warchocki S, Wright EM, Allred AF, Ahlstrom C, Manuel CS, Stasiewicz MJ, Burrell A, Roof S, Strawn LK, Fortes E, Nightingale KK, Kephart D, Wiedmann M. 2014. *Listeria floridensis* sp. nov., *Listeria aquatica* sp. nov., *Listeria cornellensis* sp. nov., *Listeria riparia* sp. nov. and *Listeria grandensis* sp. nov., from agricultural and natural environments. Int J Syst Evol Microbiol 64:1882–1889.

49. Bucur FI, Grigore-Gurgu L, Crauwels P, Riedel CU, Nicolau AI. 2018. Resistance of *Listeria monocytogenes* to Stress Conditions Encountered in Food and Food Processing Environments. Front Microbiol 9:2700.

50. Lakicevic BZ, Den Besten HMW, De Biase D. 2021. Landscape of Stress Response and Virulence Genes Among *Listeria monocytogenes* Strains. Front Microbiol 12:738470.

51. Gahan CG, Hill C. 2014. *Listeria monocytogenes*: survival and adaptation in the gastrointestinal tract. Front Cell Infect Microbiol 4:9.

52. Arcari T, Feger ML, Guerreiro DN, Wu J, O’Byrne CP. 2020. Comparative Review of the Responses of *Listeria monocytogenes* and *Escherichia col*i to Low pH Stress. Genes (Basel) 11.

53. Quereda JJ, Nahori MA, Meza-Torres J, Sachse M, Titos-Jimenez P, Gomez-Laguna J, Dussurget O, Cossart P, Pizarro-Cerda J. 2017. Listeriolysin S Is a Streptolysin S-Like Virulence Factor That Targets Exclusively Prokaryotic Cells *In Vivo*. MBio 8.

54. Rolhion N, Chassaing B, Nahori MA, de Bodt J, Moura A, Lecuit M, Dussurget O, Berard M, Marzorati M, Fehlner-Peach H, Littman DR, Gewirtz AT, Van de Wiele T, Cossart P. 2019. A *Listeria monocytogenes* Bacteriocin Can Target the Commensal Prevotella copri and Modulate Intestinal Infection. Cell Host Microbe 26:691–701 e5.

55. Hafner L, Pichon M, Burucoa C, Nusser SHA, Moura A, Garcia-Garcera M, Lecuit M. 2021. *Listeria monocytogenes* faecal carriage is common and depends on the gut microbiota. Nat Commun 12:6826.

56. Kaur G, Malik RK, Mishra SK, Singh TP, Bhardwaj A, Singroha G, Vij S, Kumar N. 2011. Nisin and class IIa bacteriocin resistance among *Listeria* and other foodborne pathogens and spoilage bacteria. Microb Drug Resist 17:197–205.

57. Chen X, Schauder S, Potier N, Van Dorsselaer A, Pelczer I, Bassler BL, Hughson FM. 2002. Structural identification of a bacterial quorum-sensing signal containing boron. Nature 415:545–9.

58. Newman H, Krajnc A, Bellini D, Eyermann CJ, Boyle GA, Paterson NG, McAuley KE, Lesniak R, Gangar M, von Delft F, Brem J, Chibale K, Schofield CJ, Dowson CG. 2021. High-Throughput Crystallography Reveals Boron-Containing Inhibitors of a Penicillin-Binding Protein with Di-and Tricovalent Binding Modes. J Med Chem 64:11379–11394.

59. Nguyen L, Schultz DC, Terzyan SS, Rezaei M, Songb J, Li C, You Y, Hanigan MH. 2022. Design and evaluation of novel analogs of 2-amino-4-boronobutanoic acid (ABBA) as inhibitors of human gamma-glutamyl transpeptidase. Bioorg Med Chem 73:116986.

60. Perez M, Crespo C, Schleissner C, Rodriguez P, Zuniga P, Reyes F. 2009. Tartrolon D, a cytotoxic macrodiolide from the marine-derived actinomycete *Streptomyces sp.* MDG-04-17-069. J Nat Prod 72:2192–4.

61. Elshahawi SI, Trindade-Silva AE, Hanora A, Han AW, Flores MS, Vizzoni V, Schrago CG, Soares CA, Concepcion GP, Distel DL, Schmidt EW, Haygood MG. 2013. Boronated tartrolon antibiotic produced by symbiotic cellulose-degrading bacteria in shipworm gills. Proc Natl Acad Sci U S A 110:E295–304.

62. Vivant AL, Garmyn D, Piveteau P. 2013. *Listeria monocytogenes*, a down-to-earth pathogen. Front Cell Infect Microbiol 3:87.

63. Rodas-Suarez OR, Flores-Pedroche JF, Betancourt-Rule JM, Quinones-Ramirez EI, Vazquez-Salinas C. 2006. Occurrence and antibiotic sensitivity of *Listeria monocytogenes* strains isolated from oysters, fish, and estuarine water. Appl Environ Microbiol 72:7410–2.

64. El-Shenawy MA, El-Shenawy MA. 2006. *Listeria spp.* in the coastal environment of the Aqaba Gulf, Suez Gulf and the Red Sea. Epidemiol Infect 134:752–7.

65. Bou-m’handi N, Jacquet C, El Marrakchi A, Martin P. 2007. Phenotypic and molecular characterization of *Listeria monocytogenes* strains isolated from a marine environment in Morocco. Foodborne Pathog Dis 4:409–17.

66. Dillon RM, Patel TR. 1992. Listeria in Seafoods: A Review. J Food Prot 55:1009–1015.

67. Lachmann R, Halbedel S, Lüth S, Holzer A, Adler M, Pietzka A, Dahouk SA, Stark K, Flieger A, Kleta S, Wilking H. 2022. Invasive listeriosis outbreaks and salmon products: a genomic, epidemiological study. Emerg Microbes Infect doi:10.1080/22221751.2022.2063075:1-30.

68. Shahamat M, Seaman A, Woodbine M. 1980. Survival of *Listeria monocytogenes* in high salt concentrations. Zentralbl Bakteriol A 246:506–11.

69. Nolan DA, Chamblin DC, Troller JA. 1992. Minimal water activity levels for growth and survival of *Listeria monocytogenes* and *Listeria innocua*. Int J Food Microbiol 16:323–35.

70. Sambrook J, Fritsch EF, Maniatis T. 1989. Molecular cloning : a laboratory manual, 2nd ed. Cold Spring Harbor Laboratory Press, Cold Spring Harbor, N.Y.

71. Hamoen LW, Smits WK, de Jong A, Holsappel S, Kuipers OP. 2002. Improving the predictive value of the competence transcription factor (ComK) binding site in *Bacillus subtilis* using a genomic approach. Nucleic Acids Res 30:5517–28.

72. Monk IR, Gahan CG, Hill C. 2008. Tools for functional postgenomic analysis of *Listeria monocytogenes*. Appl Environ Microbiol 74:3921–34.

73. van den Ent F, Löwe J. 2006. RF cloning: a restriction-free method for inserting target genes into plasmids. J Biochem Biophys Methods 67:67–74.

74. Arnaud M, Chastanet A, Debarbouille M. 2004. New vector for efficient allelic replacement in naturally nontransformable, low-GC-content, gram-positive bacteria. Appl Environ Microbiol 70:6887–91.

75. Picache JA, Rose BS, Balinski A, Leaptrot KL, Sherrod SD, May JC, McLean JA. 2019. Collision cross section compendium to annotate and predict multi-omic compound identities. Chem Sci 10:983–993.

76. Toledo-Arana A, Dussurget O, Nikitas G, Sesto N, Guet-Revillet H, Balestrino D, Loh E, Gripenland J, Tiensuu T, Vaitkevicius K, Barthelemy M, Vergassola M, Nahori MA, Soubigou G, Regnault B, Coppee JY, Lecuit M, Johansson J, Cossart P. 2009. The *Listeria* transcriptional landscape from saprophytism to virulence. Nature 459:950–6.

77. Lewis PJ, Marston AL. 1999. GFP vectors for controlled expression and dual labelling of protein fusions in *Bacillus subtilis*. Gene 227:101–10.

78. Glaser P, Frangeul L, Buchrieser C, Rusniok C, Amend A, Baquero F, Berche P, Bloecker H, Brandt P, Chakraborty T, Charbit A, Chetouani F, Couve E, de Daruvar A, Dehoux P, Domann E, Dominguez-Bernal G, Duchaud E, Durant L, Dussurget O, Entian KD, Fsihi H, Garcia-del Portillo F, Garrido P, Gautier L, Goebel W, Gomez-Lopez N, Hain T, Hauf J, Jackson D, Jones LM, Kaerst U, Kreft J, Kuhn M, Kunst F, Kurapkat G, Madueno E, Maitournam A, Vicente JM, Ng E, Nedjari H, Nordsiek G, Novella S, de Pablos B, Perez-Diaz JC, Purcell R, Remmel B, Rose M, Schlueter T, Simoes N, et al. 2001. Comparative genomics of *Listeria* species. Science 294:849–52.

